# Neuroscience-Inspired Deep Learning Brain-Machine Interface Decoder

**DOI:** 10.64898/2026.02.07.703641

**Authors:** Hong-Yun Ou, Takahiro Hasegawa, Osamu Fukayama, Eizo Miyashita

## Abstract

Brain–machine interfaces (BMIs) aim to decode motor intentions from neural activity to enable direct control of external devices. However, most existing decoders rely on monolithic architectures that fail to capture the distinct neural representations of different joint movement directions, limiting their generalizability. In this work, we propose a Single-Direction CNN-LSTM decoder inspired by motor cortex encoding mechanisms, which separately models extension and flexion dynamics through parallel CNN-LSTM branches. Each branch extracts spatial–temporal features from neural spike data and predicts directional joint variables, which are then combined by subtraction to yield the net angular velocity and torque of upper-limb joints. Using invasive recordings from a macaque during a 2D center-out reaching task, we demonstrate that our decoder achieves comparable performance to a conventional CNN-LSTM when trained on all tasks, while significantly outperforming both CNN-LSTM and linear regression baselines in cross-target generalization scenarios. Moreover, the model can capture physiologically meaningful co-contraction patterns, providing richer insights into motor control. These results suggest that incorporating neuroscience-inspired modular decoding into deep neural architectures enhances robustness and adaptability across tasks, offering a promising pathway for BMI applications in prosthetics and rehabilitation.

## 1. Introduction

Brain-machine interface is a technology that builds a bridge between the brain and the external device. A brain-machine interface (BMI or BCI) system will decode the user’s movement or selection from neural signals so the user can control external devices by brain activity.[1–3] According to the place of implanted electrodes or signals that the decoder uses, BMI devices can be divided into invasive and non-invasive [4]. Non-invasive BMIs usually use signals like EEG to estimate subjects’ imaginary movement [5,6] or sensory stimulation[7,8]. But EEG signals are always noisy and full of artifacts because the electrodes are attached to the surface of the scalp[9–11]. In contrast, for invasive signals like ECoG, LFP, or spike data, electrodes are implanted on the surface or inside the cortex, so their signal quality is much higher than EEG[12] and more explainable.

To decode information from neural signals, numerous studies have employed the population vector[13] as a feature representation. State discrimination has often been performed using linear discriminators, and solutions for linear state-space representation models have been explored through methods such as Kalman filtering[14]. Due to the non-stationary and non-Gaussian nature of neural signals, these decoders can roughly estimate the presence or absence of movement or torque, but they are not precise, and their reproducibility is lacking. With the development of artificial intelligence (AI), deep neural networks (DNN) have made remarkable achievements in non-invasive BMIs[15], and some benchmark decoder models have been proposed like EEGnet[16] and DeepConvNet[17]. Those deep learning based methods have significantly improved decoding capabilities[18]. There are also more and more researchers using DNN models to decode information from cortical neural signals and have obtained some exciting results in motor, speech[19], and cognitive function reconstruction[20]. Furthermore, by the use of data-driven feature extraction modules, researchers can get some abstract but important features of neural activity which is imperceptible for human[21,22]. In motor decoder, for example, Xie *et al*[23] predicted continuous flexion and extension of five fingers using an end-to-end DNN model with four spatial/temporal convolutional layers (CNN) as feature extractors and one LSTM layer to predict fingers activation from ECoG signals. The coefficient of determination of their model can reach 74%. Sliwowski *et al*[21] employed a CNN layer combined with several other architectures to reconstruct 3-D hand movements from the ECoG signals of a tetraplegic patient, they reported that a CNN-LSTM model with multi–time-step trajectory prediction achieved an average *R*^2^ of approximately 0.232 for decoding 3-D hand positions during a motor imagery task, representing about a 60% improvement over other architectures. Although DNN decoders were tested with higher performance in offline decoding, but their generalizability across subjects and tasks is a big challenge.

Brain science has focused on understanding the representation or encoding of Brain. Many researches neural signal decoding engineering have estimated the output movement variables in the Cartesian coordinates[21]. However, one research argued that neural activity in the motor cortex represents information in joint coordinates (eg, joint angle or joint torque) but not in Cartesian coordinates[24]. By the way, joint coordinates are essential for some special scenarios, such as prostheses and motor rehabilitation [25], in which an accurate output joint torque will give patients positive support. In our previous research, we found that the motor cortex of the monkey encoded the movement variables of the shoulder and elbow in four separate modules, representing different rotation directions of the two joints respectively-shoulder extension, shoulder flexion, elbow extension, and elbow flexion[26]. What’s more, a recent research from Tian *et al*[27] indicated that the features representing variables in different dimensions should be orthogonal. All of these evidences imply that the joint movement variables in different rotations could be encoded in different neural patterns in the motor cortex, which is not taken into account for the ordinary single backbone decoder.

Drawing on these insights, we propose a DNN architecture to separately decode the upper-limb joint movement variables, inspired by the motor cortex encoding mechanism we previously identified. Our backbone decoder is a CNN-LSTM model, which we term the Single-Direction CNN-LSTM. The “Single-Direction” design is intended to extract independent representations of joint movement variables along different directions, thereby enhancing generalization across tasks. The decoder consists of three main components: (1) independent branches that estimate extension and flexion variables separately; (2) a feature extraction module with convolutional layers to capture both spatial and temporal patterns of neural activity; and (3) an output module composed of an LSTM and fully connected (FC) layer to model the dynamic patterns, with parameters that can be fine-tuned for efficient few-shot adaptation to new tasks.

To validate the effectiveness of our approach, we employed a dataset that we recorded before from a Japanese macaque motor cortex in a 2D center-out reaching experiments shown in Figure 1 f)[26]. During the experiment, we recorded the hand position on the horizontal workspace of the monkey and then calculated the joint angle, angular velocity joint torque with the motion equation of the monkey’s upper limb. We first experimentally demonstrated the practical feasibility of our decoder in a single task and motion equation by training and testing with all the neural spike activity and movement variables we recorded. Subsequently, we evaluated the generation capability of our Single-Direction decoder. We use data from only two targets to train our decoder, then fine-tune the parameters of the output layer to mimic multi-task scenarios. We use a conventional CNN-LSTM model without branches and a simple linear regression (LR) model to give a comparison. The result shows that our Single-Direction decoder has the highest determination coefficient (*R*^2^) with competitive performance to the conventional CNN-LSTM model in single task. The main contributions of this work include:

**Figure 1.**
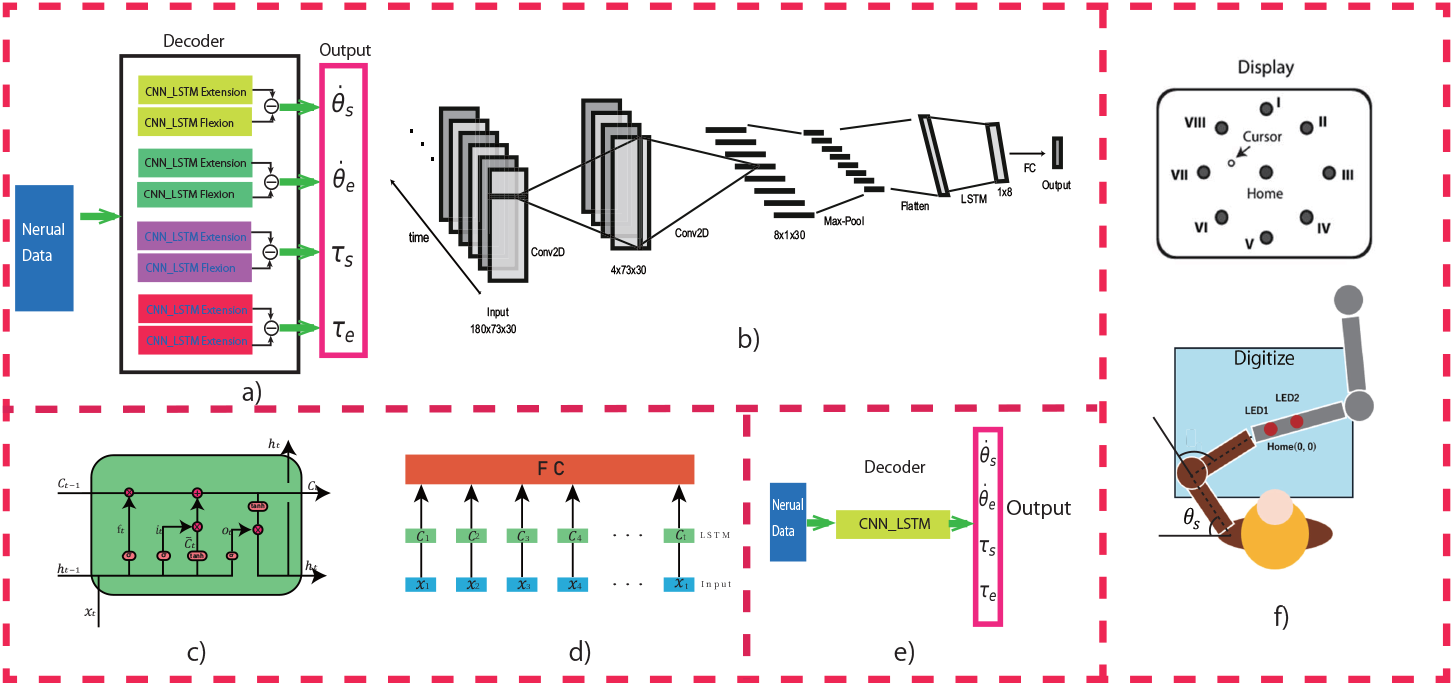
Schematic of our method. a) Structure of our single-direction model. There are multiple CNN-LSTM branches with the same structure inside the decoder. Each branch will independently estimate the variables of joints in different directions. Finally, by the subtraction between variables in counter-wise directions, the final output or net output estimated by the decoder can be given. b) Structure of CNN-LSTM branch. There are two convolution layers (CNN) to extract the temporal and spatial features from the input neural signal, respectively. Then the features will flow into an LSTM layer to estimate the dynamic sequences. Finally, a full-connected layer (FC) will do dimensional alignment to transform the sequences into the output variable in a single direction. c) Structure of the LSTM cell. d) Connection between the LSTM layer and the FC. The weights of CNNs and FC are shared in each time bin. Every hidden state of the LSTM will be used as a part of the dynamic sequence. e) Figure of the conventional CNN-LSTM decoder. f) Sketch of monkey center-out experiment and the definition of the angle of shoulder (*θ*_*s*_) and elbow (*θ*_*e*_). The rotation that makes the joint angle larger was defined as extension, while the opposite rotation was flexion. The two LEDs indicated the position of the monkey’s left hand.

1. We propose a DNN-based decoder (CNN-LSTM) for motor decoding, offering a novel formulation that bridges neuroscience mechanisms with deep learning approaches for prosthetic and rehabilitation applications.
2. We introduce the Single-Direction CNN-LSTM, which decodes joint variables independently across directions, thereby improving task-level generalizability.

## 2. Materials and Methods

### 2.1. Data preparation

Parts of dataset in our previous work[26] were used in the present study. To explain it, Neural activity in the motor cortex of Japanese monkeys (Macaca fuscata) was systematically recorded during a center-out hitting task toward peripheral 8 targets with 1-mm grids through a glass-coated Elgiloy electrode. The schematic of the experiment is shown in Figure 1 f). Instead of being recorded simultaneously with a multi-channel electrode array, the electrode recording sites were changed daily. We resampled the dataset as the monkey executed similar left-hand movement trajectories toward each target.

After spike sorting, we selected neurons with marked different preferred direction vectors (PV)[28] in flexion and extension rotation of the elbow and shoulder, which we used for our decoding. The total number of neurons we used was 73. Data of spike-firing timing was aligned with movement onset. Trial-averaged spike-firing rates for eight different targets were calculated using a 1-ms interval. Finally, the processed data from each experimental day were combined to form a large dataset.

To ensure that the monkey’s arm movements followed approximately the same trajectory toward each target across recording sessions, we first computed a reference hand trajectory using all available motion data and discarded trials that exhibited large deviations from this reference. Specifically, for each trial we calculated the angle of hand trajectory at the point located 47 mm away from the home position and clustered the trials based on these angles into three groups by K-means algorithm. The cluster containing the largest number of trials was identified as the reference, and its average trajectory was denoted as 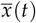 and 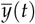 for the two spatial dimensions. Movement onset was aligned across trials using the mean onset timing, which was defined as the time point when the hand’s tangential acceleration first reached or exceeded 0.8 m/s^2^. To quantify the similarity between each individual trial and the reference, we evaluated both the trajectory error and the coefficient of determination, calculated as follows:

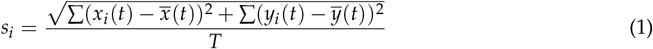

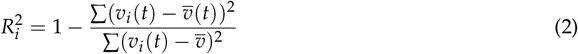

where *s*_*i*_ and 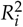 denote the trajectory error and the coefficient of determination between the hand trajectory of trial *i* and the reference. Here, *v*_*i*_(*t*) and 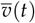 represent the hand velocities of trial *i* and the reference trajectory, respectively, and 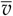 is the temporal mean of 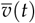. *T* denotes the total number of time steps, which in this study was defined as a 1500-ms window from the onset to termination, sampled at 1 kHz. Only trials satisfying *s*_*i*_ < 20 and 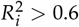 > 0.6 were retained for further analysis.

### 2.2. Data Preprocessing

Recorded neural activity was digitized at a sampling rate of 40 kHz. To extract a single neuron’s activity from the timeseries electric potential data acquired using extracellular recording, we sorted spikes using Wave-Clus (University of Leicester, Leicestershire, England). Spike activity was detected when the amplitude exceeded or fell below a threshold level, which was spike-firing timing, and data of 0.25 ms before and 0.75 ms after this timing were treated as a spike. We computed the firing rate as the inverse number of the inter-spike interval of the spikes immediately before and after the 1ms time-bin and smoothed it with a 4th-order Butterworth filter with a cutoff frequency of 7 Hz. Data of spike-firing timing was aligned upon movement onset, which was defined in this paper as the time when tangential acceleration of the hand was equal to or greater than 0.8 *m*/*s*^2^. Trial-averaged spike-firing rates for eight different targets were calculated using a 1-ms interval. After applying the moving average with a 40-ms time window, the data were resampled at a 10-ms interval.

On the one hand, the positions of the recorded LEDs were sampled at 1 kHz and converted into the monkey’s hand position in Cartesian coordinates. From these trajectories, hand velocity and acceleration were derived. The kinematic data were then down-sampled to 100 Hz. On the other hand, the shoulder and elbow joint angles, as well as their angular velocities, were computed:

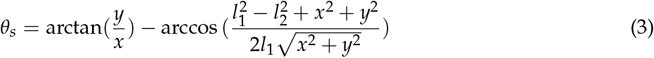

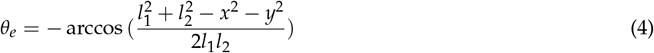

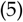

where x and y are the x- and y-axes elements of the hand position in Cartesian coordinates, and *l*_1_ and *l*_2_ are the upper arm and forearm lengths, respectively. Finally, the same filter as that used for the spike data was used to obtain the smooth hand position, velocity, joint angle, and angular velocity. All the processing was done with Matlab 2024b.

Further, the joint torques were calculated according to the arm motion equation:

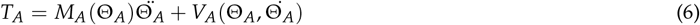

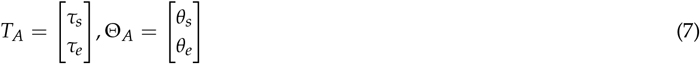

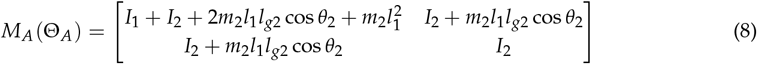

In this paper, we used four variables, shoulder angle velocity 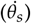, elbow angle velocity 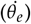, shoulder torque (*τ*_*S*_), elbow torque (*τ*_*e*_) as our outputs. *I*_1_ and *I*_2_ are inertia moments of the upper arm around the shoulder joint and the forearm around the elbow joint, respectively. *m*_2_ and *l*_*g*2_ respect to the weight of the forearm and distance from the elbow joint to the center of gravity of the forearm. *V*_*A*_ corresponds to Coriolis and centrifugal forces. In order to get the final real torque produced by joint rotation, it’s alse need to add the torque produced by pushing the manipulandum to *T*_*A*_:

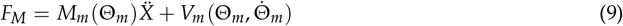

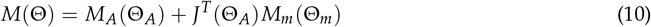

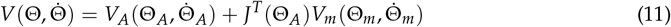

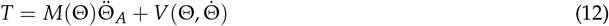

where T is the total torque vector that we use in this study, *J*^*T*^ (Θ_*A*_) is the transpose Jacobian of the relationship between joint angles and manipulandum position. The outputs movement last for 1800ms from 300ms before the onset of reaching phase towards the target to the end of the movement after hitting the target.

Before using the spike-firing data and motion data in the decoder, we divided the dataset into the train set and the test set. For the DNN decoder, to avoid overfitting, we split 80% trials (about 896 trials, each target repeated 112 times) as the train set, 10% trials validation set (about 224 trials, each target repeated 28 times), and 10% of the trials as the test set[29]. For the linear decoder, we split 80% trials as training data while the other 20% trials as test because validation is not necessary for the linear decoder. Then the spike-firing data were normalized by z-score normalization as (13) to all units[29][30], where *σ* means the standard deviation among units. The sliding window with a length of 300ms (200ms before the motion and 100ms after [31]) cleaved the normalized spike data into segments with the same number of sampling points as the motion data. Finally, the input data of DNN decoder has the shape of [*B* × *T* × *C* × *L*], in which *B, T, C, L* represent batch size, motion time, number of units, and the length of the sliding window, respectively. For the linear decoder, we just concatenate the spike data along the temporal dimension, so the input shape of the linear decoder is [(*B* · *T*) × *C*][32].

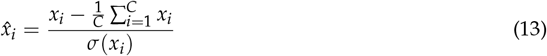

### 2.3. Decoder Model

In this study, we employed a deep neural network (DNN) model consisting of multiple CNN-LSTM branches to decode the angular velocity and torque of the monkey’s shoulder and elbow joints. The architecture of the decoder is illustrated in Figure 1a. The decoder comprises four pairs of CNN-LSTM branches, each designed to independently estimate the parameters for extension and flexion movements corresponding to a single output. Since the final output—the net joint parameter is obtained by subtracting the flexion-related component from the extension-related one[33], a subtraction layer is incorporated after the two branches. Given that our decoder independently estimates extension and flexion parameters, we refer to it as the single-direction decoder. The internal structure of each CNN-LSTM branch is shown in Figure 1b, and the detailed configuration is described below.

#### 2.3.1. Convolutional layers

Similar to EEGnet[16], we employed two convolutional layers to extract temporal and spatial features from the spike data independently. The first separable convolution layer focuses on temporal feature extraction, capturing the specific time intervals that the network is most responsive to—referred to as the “lag time”[34][35]. The temporally filtered data is then passed to a second separable convolutional layer, which is responsible for extracting spatial features. The learned weights in this layer indicate which neuronal units the model attends to most, i.e., those whose activity exhibits the highest correlation with the target output. The output from the spatial convolution layer is subsequently flattened and passed through a max-pooling operation, after which it is fed into an LSTM layer for temporal sequence modeling.

#### 2.3.2. LSTM layer

LSTM is a special form of recurrent neural network (RNN) invented by Hochreiter and Schmidhuber in 1997[36]. When the time sequence is very long, traditional RNN faces the problem of gradient vanishing or explosion[37]. LSTM solves this problem by using multiple gates to control the data flow. Since its appearance, LSTM has been widely used in many time sequence prediction problems. There are also many researches who use LSTM as a decoder in intracranial BMI and have obtained some exciting results. The equation and structure of LSTM cell is shown as (13)-(18) and Figure 1 c), where *f*_*t*_, *i*_*t*_, *O*_*t*_ mean forget gate, input gate, and output gate at time *t* respectively. *x*_*t*_, *h*_*t*_, *C*_*t*_ mean input data, hidden state and statement of memory cell at time *t*. Detailed information is shown in [36]. In our decoder, we used an LSTM layer to fit the dynamic sequences of the output parameters. Connected with a readout full-connected layer, these two layers can do as the output layer of our decoder, as in Figure 1 d). Since the angular velocity and torque during extension or flexion are always non-negative, we employ the ReLU function [38] as the activation function of the fully connected layer.

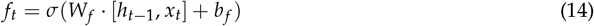

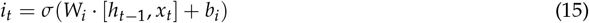

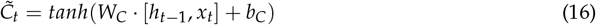

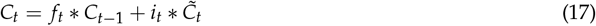

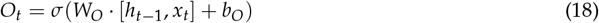

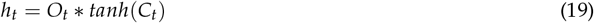

### 2.4. Single-Direction CNN-LSTM decoder

As shown in Figure 1, there is a pair of branches inside the model for each output variable in the Single-Direction CNN-LSTM decoder. First, two convolution layers will extract the spatial and temporal features from the input neural activity of orthogonal extension and flexion variables independently:

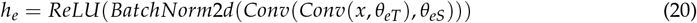

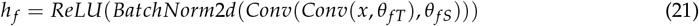

where *h*_*e*_ and *h* _*f*_ represent extension and flexion, respectively. Conv corresponds to the two convolution layers, *θ*_*eT*_ and *θ*_*eS*_ denote the learnable parameters inside these two layers that extract spatial and temporal features, respectively.*BatchNorm*2*d* represents batch normalization for stable training, and *ReLU*(·) is Rectified Linear Unit. Then the extracted spatial-temporal features will flow into the output layer, which is made up of LSTM and Fully connected layer, to fit the dynamics of variables and output the extension and flexion estimation:

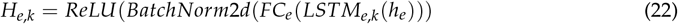

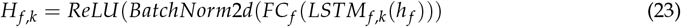

where *H*_*e,k*_ and *H*_*f*,*k*_ denote the outputs of the extension and flexion branches at the *k*-th time step, respectively. *FC*_*e*_ and *FC*_*f*_ represent the fully connected layers of the extension and flexion branches. Since the outputs of ReLU are always non-negative, the estimated values for extension and flexion are also non-negative. Finally, the net variables are obtained by subtracting the flexion branch output from the extension branch output:

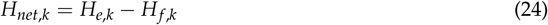

### 2.5. Conventional CNN-LSTM decoder

We compared our single-direction decoder with the conventional CNN-LSTM decoder without branches[23]. The structure of the conventional CNN-LSTM decoder is shown in Figure 1 e). A single CNN-LSTM will output all the parameters (angle velocities and torques) at the same time. In order to speed up the training and keep the training balance between different outputs in case of multiple regression outputs, we need to scale the output series, so we also do the z-score normalization to the output motion data as input spike data.[39]

### 2.6. Linear decoder

To give a more complete comparison, we also compare the performance between our single-direction decoder and the linear decoder[26][32]. The model of the linear decoder is the same as the equation (25)(26), where X means the input spike data with the shape [(*B* · *T*) × *C*] that has been explained before, P means the output parameter with the shape [(*B* · *T*) × 4]. We first calculated *β* based on the training dataset and then tested the trained decoder on the test set.

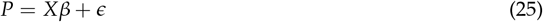

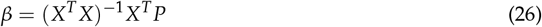

### 2.7. Fine-tuned and generalizability test

In center-out experiments, motion toward a target in a different direction can be classified as a different sub-task, as the dynamics in each target have different patterns and initial states[40]. So we used some targets to train the decoder and tested on the other targets to validate the generalizability of the decoder.

First, we plot the motion space of each target using the elbow and shoulder variables as horizontal and vertical coordinates as shown in Figure 2. From the figure, we chose target IV and target V to train our decoder because the trajectories of these two targets cover the largest area of the motion space with the fewest trajectories. So we think the decoder trained by these two targets has enough knowledge to estimate the parameters of other decoders.

**Figure 2.**
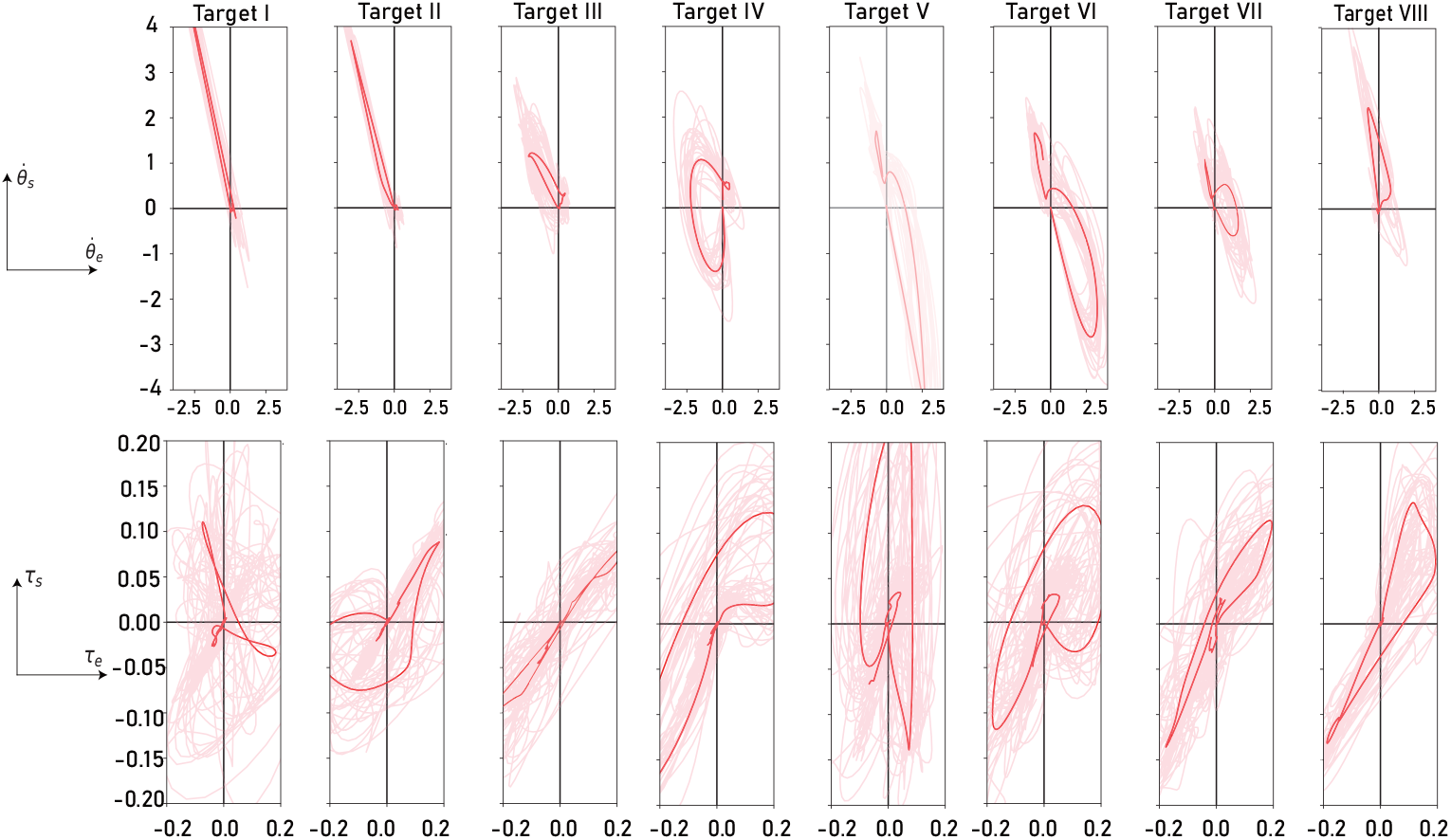
Motion space of all the eight targets. Pink shallow lines represent trajectories of each trial, while the red line represents the mean trajectory of all trials in each target. Upper row: motion space of angle velocities; Lower row: motion space of joint torques

After training the decoder, we fine-tuned its output layer—including the LSTM layer and the connected fully connected (FC) layer—using a small number of trials from a target with a different movement direction. The fine-tuned decoder was then evaluated on the remaining data from this target[41]. During fine-tuning, the convolutional layers were kept fixed, based on the assumption that they extract general features shared across all directions, while the temporal dynamics captured by the output layers vary depending on the target direction.

### 2.8. Environment and hyperparameter

The experiments were conducted on a Windows PC running Python 3.10 and TensorFlow 2.10, with an RTX A4000 GPU (16GB memory). The hyperparameter settings are summarized in Section 3.2. To mitigate potential bias from data ordering, all training and testing procedures were performed using 5-fold cross-validation. The learn rate was set to 1.0 × 10^*−*3^ for training and 1.0 × 10^*−*4^ for fine-tuning. For each target—except for Target IV and Target V—35% of the trials (50 trials) were used for fine-tuning the decoder, while the remaining trials were used for testing. We chose the mean squared error (MSE) as the loss function and used Adam[42] to optimize the training parameters. Early stopping with a patience of 20 epochs was applied during both training and fine-tuning to prevent overfitting. All data splits in this study were performed randomly. The batch size was set to 32, and the dropout rate was 0.5.

We used the coefficient of determination, *R*^2^, as a measure of the strength of the linear association between the predicted and the ground-truth kinematic parameters[43]. The definition of *R*^2^ is:

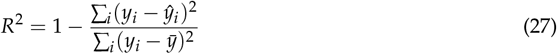

where *y*_*i*_ and 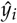 are the ground truth and prediction. 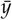 is the average of truth. The larger the *R*^2^ is, the better the performance.

## 3. Results

In this study, we developed a deep neural network architecture termed the Single-Direction CNN-LSTM Decoder, specifically designed to decode the angular velocity and joint torque of a monkey’s upper limb with isolated extension or flexion movements. The decoder incorporates parallel CNN-LSTM branches, each of which independently learns spatial and temporal representations corresponding to either extension or flexion from the input neural signals. These learned features are then propagated through an output module composed of an LSTM layer followed by a fully connected (FC) layer, which maps the high-dimensional feature representations to continuous dynamic output sequences. We hypothesize that this modular structure, by explicitly separating the directional dynamics, enhances the model’s capacity for disentangling complex motor representations and thus offers improved generalizability across movement contexts, compared to conventional monolithic decoding architectures.

To evaluate the model’s decoding performance, we utilized comprehensive neural spike-firing and kinematic datasets recorded during center-out reaching tasks in a non-human primate. For assessing generalizability, the model was trained exclusively on trials involving Targets 4 and 5, and then evaluated on trials targeting other directions. This cross-target evaluation was designed to mimic transfer learning scenarios. We benchmarked our model against a conventional CNN-LSTM decoder and a baseline linear regression model. Quantitative comparisons demonstrated that our Single-Direction CNN-LSTM Decoder achieved superior generalization performance, highlighting its potential for robust neural decoding in variable motor tasks.

### 3.1. Validation on all data

The decoding performance across all targets is shown in Figure 3. Panel e) presents the *R*^2^ values for shoulder and elbow angular velocity and torque decoded by the Single-Direction CNN-LSTM model, the conventional CNN-LSTM model, and the Linear Regression (LR) model. The two deep learning-based decoders significantly outperformed the LR model, with improvements of approximately 48%, 41%, 59%, and 76% for shoulder angular velocity, elbow angular velocity, shoulder torque, and elbow torque, respectively (five-fold cross validation, *p *<* 0.05, independent t-test).

**Figure 3.**
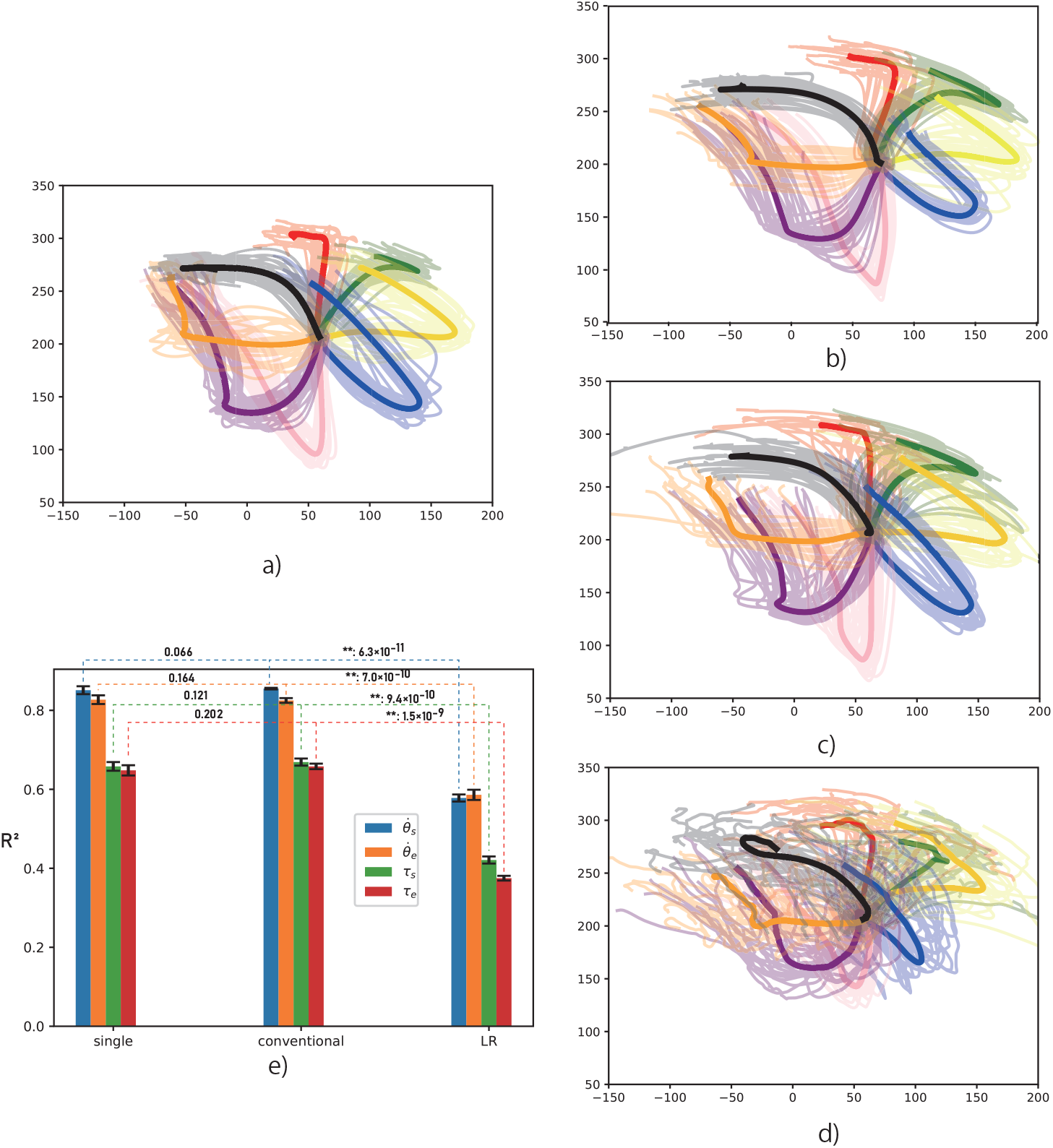
Trajectories for all eight targets based on angular velocities estimated by three decoders. Solid lines represent the average trajectory per target; faint lines represent individual trial trajectories. Different color represent trajectories of different target. a) Ground-truth trajectories recorded during the experiment; b) Trajectories reconstructed by the Single-Direction model; c) conventional CNN-LSTM model; d) Linear Regression (LR) model; e) *R*^2^ scores for each output parameter across the three decoders. single: single-direction CNN-LSTM model; conventional: conventional CNN-LSTM model; numbers on the line: p-values, independent t-test, **:p *<* 0.01; 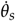: shoulder angular velocity; 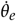: elbow angular velocity; *τ*_*s*_: shoulder torque; *τ*_*e*_: elbow torque.

Hand trajectories reconstructed from shoulder and elbow angular velocities are shown in panels (b)–(d). These results further demonstrate the poor performance of the LR decoder across all targets. Although the average trajectory for each target is relatively clear in the LR decoder (x: *R*^2^ = 0.578; y: *R*^2^= 0.586), the trial-by-trial trajectories are highly scattered and difficult to distinguish, especially when compared to the more consistent outputs from the two neural network decoders. This suggests that the LR model lacks stability at the single-trial level.

However, the *R*^2^ scores of the Single-Direction decoder did not differ significantly from those of the conventional CNN-LSTM decoder trained on all eight targets (p = 0.066, 0.164, 0.121, and 0.202 for shoulder angular velocity, elbow angular velocity, shoulder torque, and elbow torque, respectively).

The mean *R*^2^ values across five-fold cross-validation for both CNN-LSTM models were approximately 0.825–0.855 for angular velocity and 0.650–0.700 for torque.

### 3.2. Generlizability

In this experiment, we first used all the data from Target IV and Target V to pre-train a decoder. We then fine-tuned the decoder for each of the other targets individually, using 40% of their data (approximately 56 trials per target). Finally, we evaluated the fine-tuned decoder on the remaining data. Each fine-tuning process was conducted for 100 epochs. The test results for the six other targets are presented in Figure 4.

**Figure 4.**
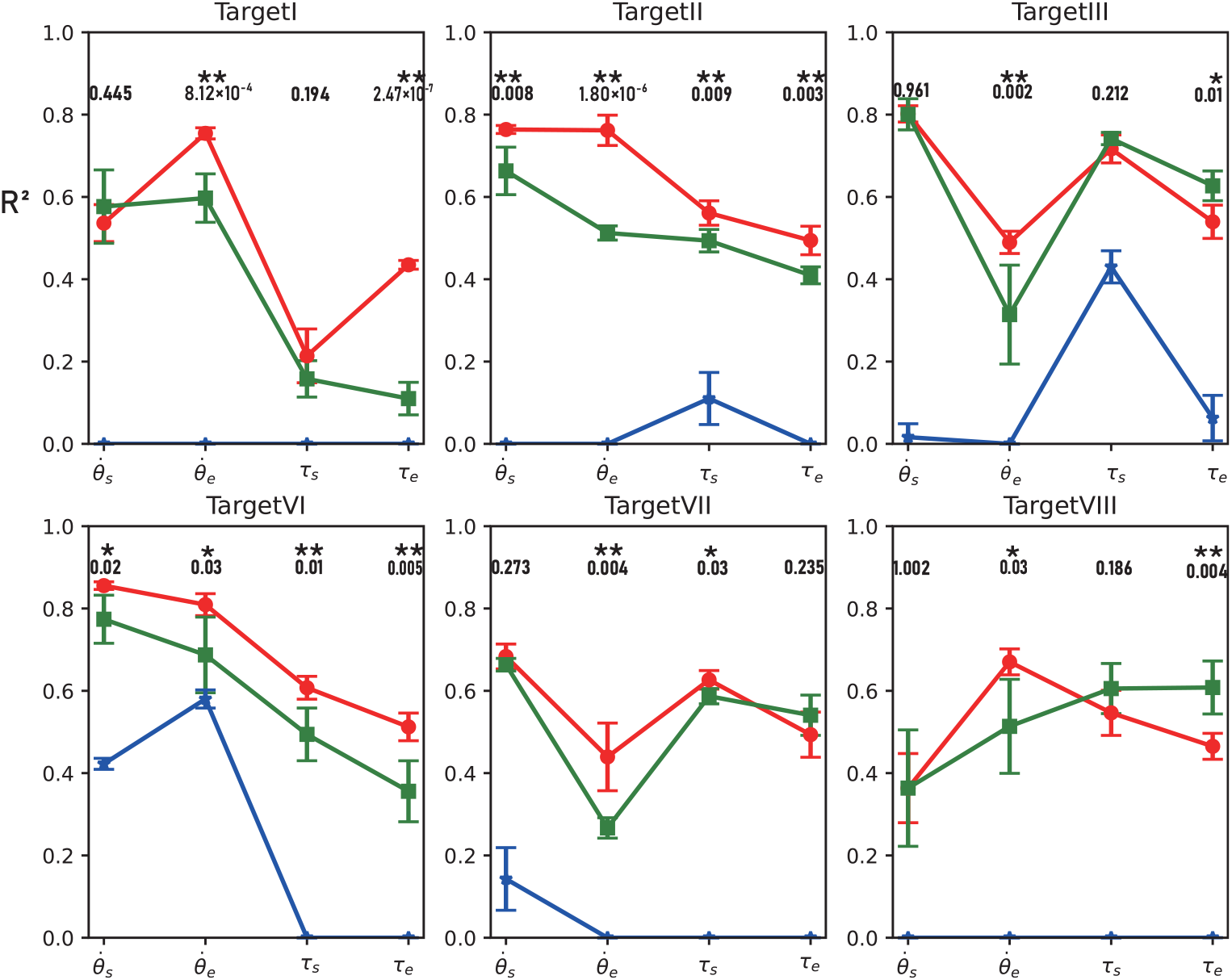
Results of the generalizability test for each target. Each colored point represents the mean *R*^2^ value for a given model, with error bars indicating one standard deviation from five-fold cross-validation. Negative *R*^2^ values were set to zero. Bold numbers in the figure represent the p-values; 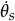: shoulder angle velocity; 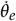: elbow angle velocity; Red: Single-Direction decoder; Green: Conventional CNN-LSTM decoder; Blue: Linear regression decoder. *τ*_*s*_: shoulder torque; *τ*_*e*_: elbow torque. *:p *<* 0.05, **:p *<* 0.01, independent t-test.

From the figure, it is evident that the deep learning–based decoder achieved strong performance across most output parameters for each target. Although the Single-Direction decoder we developed showed only minor differences from the conventional CNN-LSTM model when validated on the full dataset, the fine-tuned model for each target clearly outperformed the conventional CNN-LSTM decoder in most cases (five-fold cross-validation, p *<* 0.05, independent t-test), particularly for Target II and Target VI. For Target II, the mean *R*^2^ values of the fine-tuned model were approximately 15.1%, 48.7%, 13.6%, and 20.7% higher than those of the conventional CNN-LSTM decoder for shoulder angular velocity, elbow angular velocity, shoulder torque, and elbow torque, respectively. For Target VI, the corresponding improvements were 10.6%, 17.8%, 22.9%, and 43.9%.

**Table 1.** Hyperparameter.

| Model | Layer | Hyperparameter | Shape of Output |
| --- | --- | --- | --- |
| Extension | Input | (batch, 180, 73, 30) | (batch, 180, 73, 1) |
| | Conv_2D | kernel: $(1, 30) \times 4$, activation: ReLU | (batch, 180, 73, 1, 4) |
| | Conv_2D | kernel: $(73, 1) \times 2$, activation: ReLU | (batch, 180, 1, 30, 2) |
| | Pooling2D | pool size: $(1, 4)$, type: average | (batch, 180, 1, 27, 2) |
|  | Flatten | – | (batch, 180, 54) |
|  | LSTM | units: 8, activation: tanh | (batch, 180, 8) |
|  | FC | outputs: 1, activation: ReLU | (batch, 180, 1) |
| Flexion | Input | (batch, 180, 73, 30) | (batch, 180, 73, 1) |
| | Conv_2D | kernel: $(1, 30) \times 4$, activation: ReLU | (batch, 180, 73, 1, 4) |
| | Conv_2D | kernel: $(73, 1) \times 2$, activation: ReLU | (batch, 180, 1, 30, 2) |
| | Pooling2D | pool size: $(1, 4)$, type: average | (batch, 180, 1, 27, 2) |
|  | Flatten | None | (batch, 180, 54) |
|  | LSTM | units: 8, activation: tanh | (batch, 180, 8) |
|  | FC | outputs: 1, activation: ReLU | (batch, 180, 1) |
| Net | Subtraction | None (Extension – Flexion) | (batch, 180, 1) |
|  | Concatenation | None | (batch, 180, 4) |
| Conventional | Input | (batch, 180, 73, 30) | (batch, 180, 73, 1) |
| | Conv_2D | kernel: $(1, 30) \times 4$, activation: ReLU | (batch, 180, 73, 1, 4) |
| | Conv_2D | kernel: $(73, 1) \times 2$, activation: ReLU | (batch, 180, 1, 30, 2) |
| | Pooling2D | pool size: $(1, 4)$, type: average | (batch, 180, 1, 27, 2) |
|  | Flatten | None | (batch, 180, 54) |
|  | LSTM | units: 8, activation: tanh | (batch, 180, 8) |
|  | FC | outputs: 4, activation: ReLU | (batch, 180, 4) |

### 3.3. Features extracted by the decoder

To further examine the features extracted by the Single-Direction decoder, we analyzed the weights of the two CNN layers[44], which reflect spatial and temporal representations, as shown in Figure 5 and Figure 6. In Figure 5, the black diagonal line indicates equal weights in the extension and flexion directions. The results show that most units exhibit distinct weights between these two directions, and this trend varies across output variables. Specifically, for elbow torque, many units display identical or nearly identical weights in both directions, whereas for shoulder torque and shoulder angular velocity, a larger number of units tend to exhibit stronger weights in either flexion or extension. Similarly, Figure 6 illustrates that temporal weights also differ across output variables as well as between directions. These results on spatial and temporal features suggest that the encoding and decoding patterns differ across variables and directions, which is consistent with our hypothesis as well as with findings from our previous research [26].

**Figure 5.**
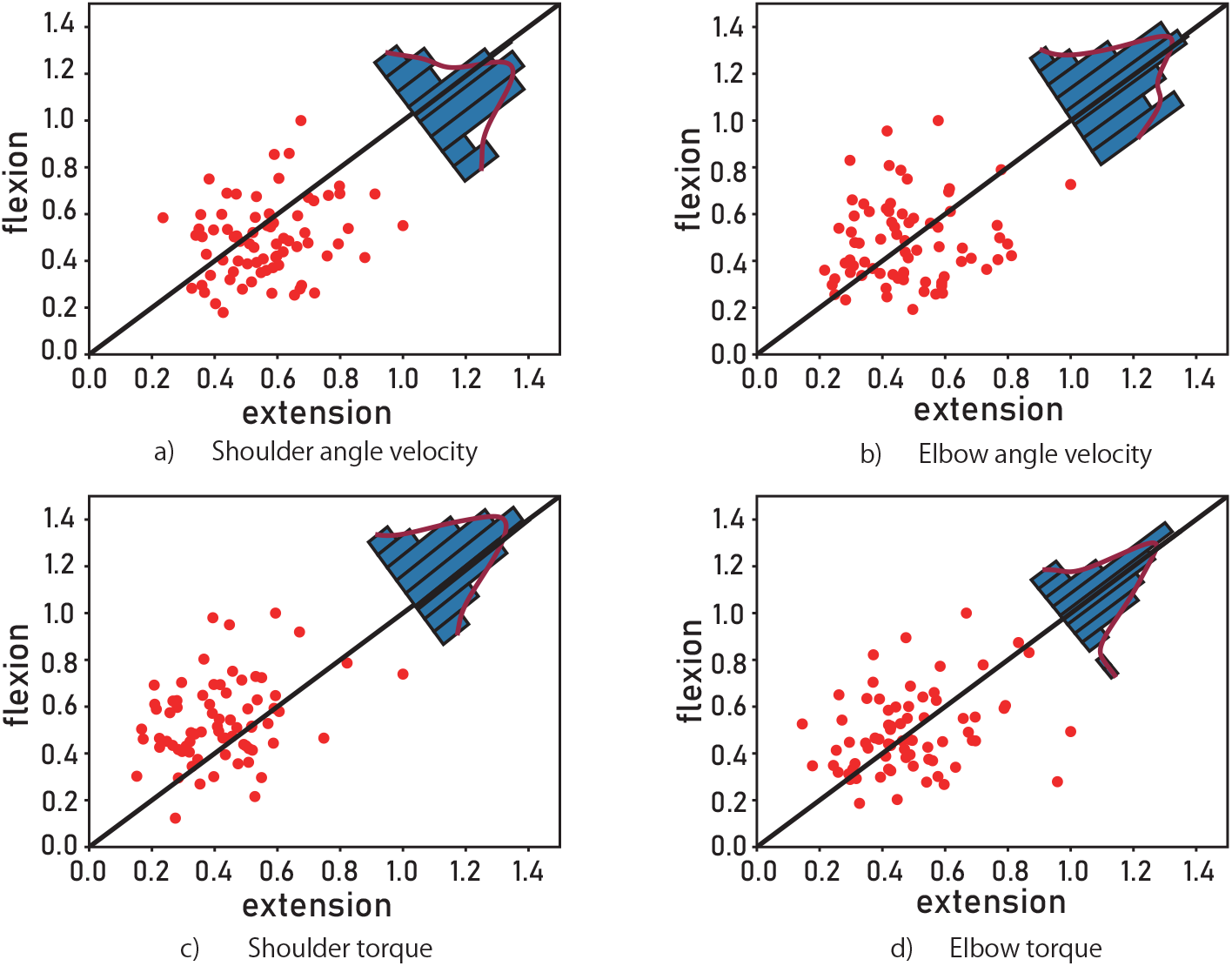
Spatial features extracted by the Single-Direction decoder. Each red point represents a unit of the input neural data, with the abscissa and ordinate indicating the unit’s weight in the extension and flexion directions, respectively. The blue histogram and overlaid red curve above the diagonal illustrate the distribution of units across different weight values (p *<* 0.05, independent t-test).

**Figure 6.**
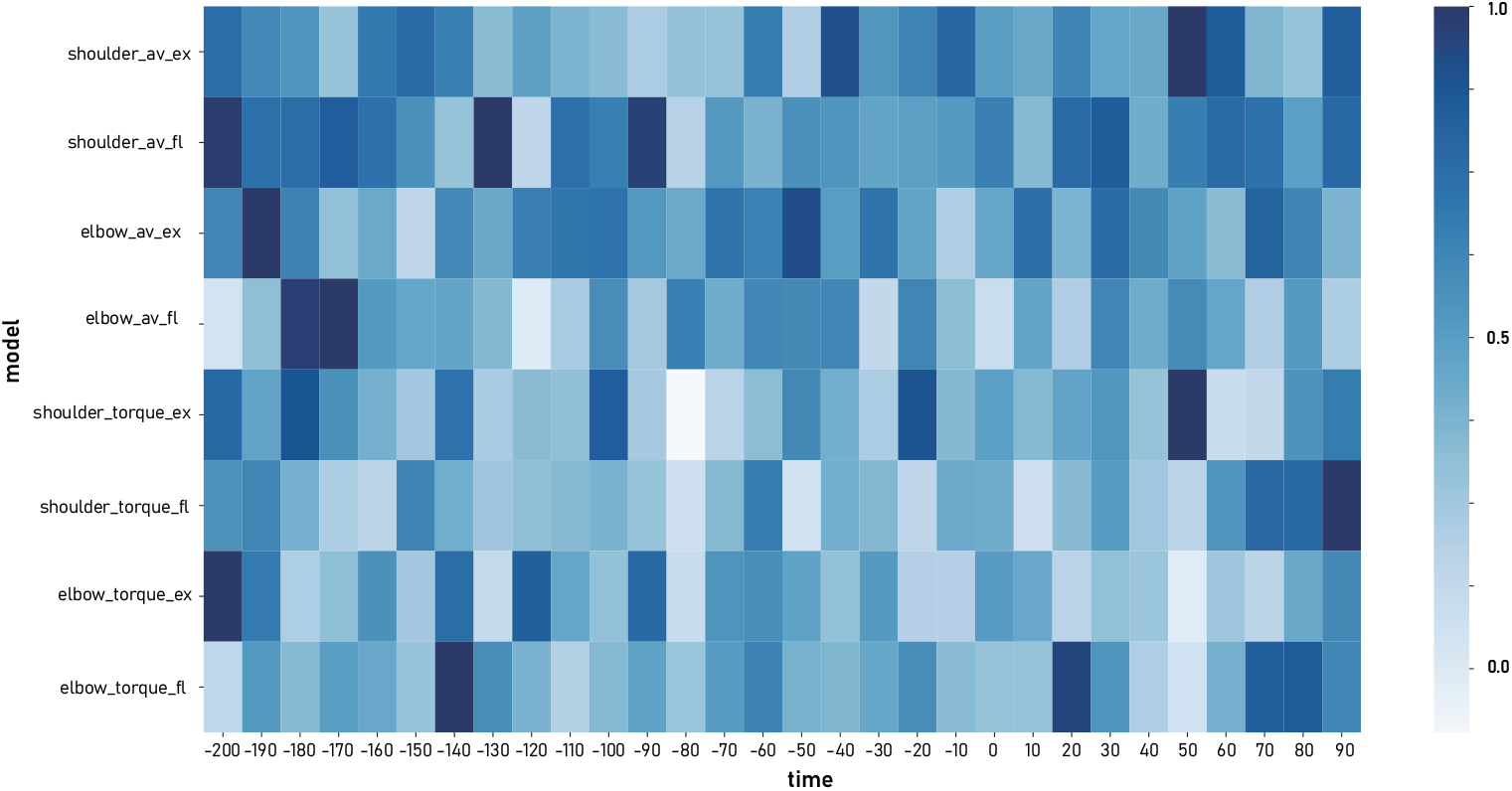
Temporal features extracted by the Single-Direction decoder. The x-axis shows time bins of the input neural data (–200 ms to 100 ms, with negative values indicating pre-movement onset), and the y-axis denotes decoder branches estimating variables in different directions. All weights are normalized to the range 0.0–1.0 within each branch.

### 3.4. Estimation of co-contraction increase the generalizability

Above results indicate that the CNN-LSTM–based Single-Direction model achieves superior generalizability compared with both the conventional CNN-LSTM decoder and the linear regression model. We attribute this advantage to the model’s ability to account for muscle co-contraction, a physiological phenomenon in which agonist and antagonist muscles are simultaneously activated. This mechanism is crucial for primates and humans to maintain upper-limb stability and adapt to different environments. For instance, when lifting a cup filled with water, the movement trajectory is largely similar to that of lifting an empty cup; however, both agonist and antagonist muscles must contract simultaneously and with comparable amplitude to achieve the motion goal while ensuring hand stability. Previous studies have shown that such co-contraction information is encoded in neural activity within the M1 region [45]. Nevertheless, conventional decoders struggle to independently extract and utilize this information.

By independently decoding motion variables in extension and flexion, the Single-Direction model can more accurately capture the nuanced neural representations underlying different movement directions, which likely contributes to its superior performance across tasks. To estimate the co-contraction of the monkey’s shoulder and elbow from the decoder output, we first extract the predicted extension and flexion torques from each branch. Since both flexion and extension variables are non-negative, we define the co-contraction torque of the shoulder and elbow as the minimum of the two, as illustrated below:

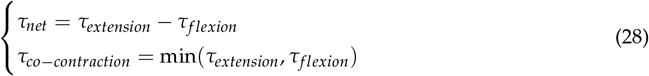

Figure 7 presents the results. The Single-Direction decoder can independently estimate extension and flexion variables, enabling the identification of periods when agonist and antagonist muscles are simultaneously activated, as indicated by the red line. Importantly, this red line—representing the co-contraction torque—peaks when the net torque equals zero, indicating that although the joint motion remains unchanged, the joint stiffness increases. Such a phenomenon cannot be captured by the conventional CNN-LSTM or linear regression decoders, as they do not separately model extension and flexion variables.

**Figure 7.**
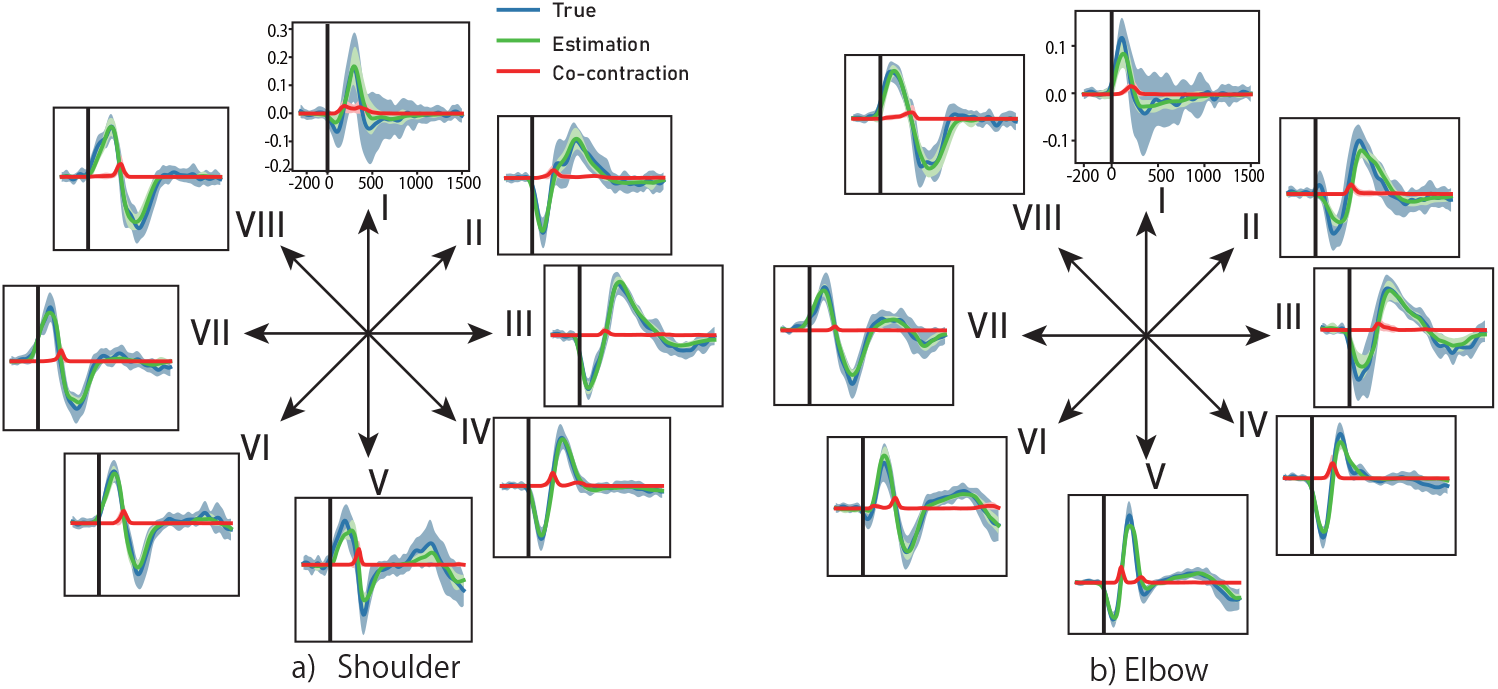
Torque results. Arrows indicate movement targets. Solid lines show trial averages, shaded areas the standard deviation. (a) Shoulder; (b) Elbow. Blue: experimental torque; Green: decoder-estimated torque; Red: decoder-estimated co-contraction torque.

## 4. Discussion

With the rapid advancement of deep learning, neural decoders have shown impressive performance. Yet, their limited generalizability across different workspaces remains a critical challenge. Our Single-Direction CNN-LSTM model, which independently estimates extension and flexion through multiple branches, addresses this issue by capturing finer motor details such as joint co-contraction. This design appears to improve adaptability compared with conventional CNN-LSTM or linear regression decoders. In comparison with previous studies that decoded joint angle variables from motor cortex activity using LSTM models and reported an average *R*^2^ of approximately 0.8 [18,23], our decoder demonstrates superior performance and is further capable of estimating joint torque simultaneously. We also observed that, for elbow torque in Target VIII, the fine-tuned conventional CNN-LSTM decoder achieved higher *R*^2^ values than the Single-Direction decoder. This improvement may stem from overfitting during fine-tuning on Target VIII, as evidenced by a decreasing training loss along with an increasing test loss. In contrast, the pre-trained linear regression decoder demonstrated limited generalizability across the other six targets.

The results confirm that the Single-Direction decoder not only outperforms linear regression but also shows advantages over the conventional CNN-LSTM in fine-tuning tests. Nevertheless, three issues merit further discussion:

### 4.1. Limitation of fine-tuning

In this study, we first pre-trained a decoder using data from Targets IV and V, and then fine-tuned the model’s output layer on the remaining targets. As a result, the model trained on only two targets was able to generalize to others. However, fine-tuning still requires sufficient data and may face the problem of overly biased pre-trained model. [46][47], which are often limited in real clinical scenarios. Currently, many approaches have been proposed to address this challenge through few-shot learning techniques, such as meta-learning [46], domain adaptation [48], and metric learning [49], which transfer knowledge across multiple tasks to a new one. Therefore, validating our Single-Direction model within these frameworks is crucial for its practical application in BMI systems.

### 4.2. Musculoskeletal model

To calculate the joint torque, we employed a musculoskeletal motion model of the monkey’s upper limb. However, accurately measuring limb length and mass is challenging, as these measurements are often subject to errors and artifacts. Consequently, obtaining precise variables for the musculoskeletal model is difficult, which in turn affects the performance of data-driven decoders.

To address this issue, one potential solution is to generate synthetic movement data based on noisy measurements to augment the training set [50]. In future work, we aim to explore data augmentation techniques to mitigate measurement inaccuracies in limb variables, thereby enhancing the overall performance of the decoder.

### 4.3. Co-contraction judgment

We operationally define the co-contraction torque as the minimum of the estimated extension and flexion torques, as expressed in (28). This definition is motivated by the following observations: (1) both extension and flexion torques are inherently non-negative, (2) their difference approximates the net torque with reasonable accuracy, and (3) the derived co-contraction tends to peak when the net torque is close to zero. Nevertheless, this measure should be regarded as a proxy rather than a definitive estimate of co-contraction. A rigorous validation would require comparison with ground-truth quantitative co-contraction data, which remains difficult to acquire[51].

## 5. Conclusion

This study demonstrates that a Single-Direction CNN-LSTM model, guided by prior neuroscience knowledge, can effectively decode upper-limb joint movements with high generalizability. Using only limited training targets, the model achieves comparable performance to conventional CNN-LSTM approaches, while its strength may lie in predicting joint co-contraction. These findings suggest that incorporating neuroscience principles into brain–machine interface design provides a promising pathway toward developing universal and robust BMI systems.

## 6. Acknowledgement

This research was supported by AMED under Grant Numbers JP23ym0126812 and JP24ym0126812.

## Disclaimer/Publisher’s Note

The statements, opinions and data contained in all publications are solely those of the individual author(s) and contributor(s) and not of MDPI and/or the editor(s). MDPI and/or the editor(s) disclaim responsibility for any injury to people or property resulting from any ideas, methods, instructions or products referred to in the content.

